# Proteomic-based stratification of intermediate-risk prostate cancer patients

**DOI:** 10.1101/2023.03.03.530910

**Authors:** Qing Zhong, Sun Rui, Adel T. Aref, Zainab Noor, Asim Anees, Yi Zhu, Natasha Lucas, Rebecca C. Poulos, Mengge Lyu, Tiansheng Zhu, Bo Wang, Guo-Bo Chen, Yingrui Wang, Xuan Ding, Dorothea Rutishauser, Niels J. Rupp, Jan H. Rueschoff, Cédric Poyet, Thomas Hermanns, Christian Fankhauser, María Rodríguez Martínez, Wenguang Shao, Marija Buljan, Janis Frederick Neumann, Andreas Beyer, Peter G. Hains, Roger R. Reddel, Phillip J. Robinson, Ruedi Aebersold, Tiannan Guo, Peter J. Wild

**Affiliations:** ProCan^®^, Children’s Medical Research Institute, Faculty of Medicine and Health, The University of Sydney, Westmead, New South Wales, Australia; Westlake Institute for Advanced Studies, Westlake University, China; Urology & Nephrology Center, Department of Urology, Clinical Research Institute, Zhejiang Provincial People’s Hospital, People’s Hospital of Hangzhou Medical College, Hangzhou, Zhejiang, China; Department of Pathology and Molecular Pathology, University Hospital Zürich, Zürich, Switzerland; Department of Urology, University Hospital Zürich, Zürich, Switzerland; Department of Urology, Cantonal Hospital Lucerne, Lucerne, Switzerland; IBM Zürich Research Laboratory, Zürich, Switzerland; State Key Laboratory of Microbial Metabolism, Joint International Research Laboratory of Metabolic and Developmental Sciences, School of Life Sciences and Biotechnology, Shanghai Jiao Tong University, Shanghai, China; Empa - Swiss Federal Laboratories for Materials Science and Technology, St. Gallen, Switzerland; Swiss Institute of Bioinformatics (SIB), Lausanne, Switzerland; CECAD, University of Cologne, Cologne, Germany; Department of Biology, Institute of Molecular Systems Biology, ETH Zürich, Switzerland; Faculty of Science, University of Zürich, Zürich, Switzerland; Dr. Senckenberg Institute of Pathology, University Hospital Frankfurt, Frankfurt am Main, Germany; Frankfurt Institute for Advanced Studies (FIAS), Frankfurt am Main, Germany

## Abstract

Gleason grading is an important prognostic indicator for prostate adenocarcinoma and is crucial for patient treatment decisions. However, intermediate-risk patients diagnosed in Gleason Grade Groups (GG) 2 and GG3 can harbour either aggressive or non-aggressive disease, resulting in under- or over-treatment of a significant number of patients. Here, we performed proteomic, differential expression, machine learning, and survival analyses for 1,348 matched tumour and benign sample runs from 278 patients. Three proteins (F5, TMEM126B and EARS2) were identified as candidate biomarkers in patients with biochemical recurrence. Multivariate Cox regression yielded 18 proteins, from which a risk score was constructed to dichotomise prostate cancer patients into low- and high-risk groups. This 18-protein signature is prognostic for the risk of biochemical recurrence and completely independent of the intermediate GG. Our results suggest that markers generated by computational proteomic profiling have the potential for clinical applications including integration into prostate cancer management.

## INTRODUCTION

Prostate Cancer (PCa) is the third most common cancer among men by incidence (14.1%) and the fifth in terms of cancer-related mortality worldwide (among men 7%)^1^. In Australia, Western Europe and North America, PCa is the most commonly diagnosed cancer among men and the second most common cause of cancer-related death^1^. PCa is a highly heterogeneous disease, and so far, most of the treatment-decision algorithms depend on risk stratification based on tumour stage, Prostate Specific Antigen (PSA) level at the time of diagnosis and the Gleason Grade Group (GG)^2^. Although this clinical risk stratification has been shown to be of prognostic and predictive value^3^, better biomarkers are still required to improve patient stratification.

The Gleason score (GS) is a grading classification of the growth pattern of prostatic adenocarcinoma. The total GS (from 6 to 10) represents the summation of the two most common predominant scores (from 1 to 5) within the specimen^4^. Despite its proven prognostic value, there was major heterogeneity within the GS7, with a differential prognosis observed between the GS7 (3+4) and GS7 (4+3) patterns^5^. Because of this, the International Society of Urological Pathology (ISUP) developed a modification to the GS system in 2014 and created a new grading of five groups, with the aim of differentiating GS7 (3+4) (termed GG2 in ISUP 2014) from GS7 (4+3) (GG3)^6^. The prognostic value of the GG system was validated in multiple cohorts, although its accuracy did not significantly differ from the older GS system^7^. In addition, for the new GG system there is controversy regarding the value of incorporating the percentage of GS4 within the GG2 and GG3, among other questions^8^. This was addressed in the ISUP 2019 modification for PCa grading, which recommends reporting the percentage of GS4 patterns in any GG2 or GG3 case^9^. Despite all of these modifications, both the GS and GG systems still have several limitations, including relatively long processing time, subjectivity, inter-observer variability, and unsatisfactory prediction of outcomes ^10,11,12,13,14,15,16^.

Therefore, there is a need to develop better prognostic biomarkers that can be interpreted either alone or when integrated with clinico-pathological features. There have been several ongoing efforts that aim to identify better molecular and genetic-based prognostic biomarkers. These include metabolomic-based biomarkers^17^, mRNA-based biomarkers such as SelectMDx^®^ and ExoDx Prostate IntelliScore^®^, urine biomarkers such as PCA3, and genetic tissue-based biomarkers such as Oncotype DX^®^, Confirm MDx^®^, Prostatype^®18,19,20^, and Prolaris^®21^. Of note, only PCA3 and Polaris^®^are FDA-approved for specific indications^21^. More recently, Proclarix showed better accuracy in detecting clinically significant PCa compared to free PSA percentage alone^22^, with its utility in clinical practice yet to be confirmed.

During the last decade, proteogenomics has revealed a range of intra-patient network effects across multi-omic layers^15^, and has described novel regulated pathways that are related to PCa progression^23^ and PCa aggressive phenotypes^24,25^. Proteogenomics appears to have the potential to provide a deep and dynamic interpretation of the underlying pathways related to cancer development, classification, and progression^26^. However, the lack of robust proteomic analyses of large cancer cohorts^27^ has limited the incorporation of proteomic-based biomarkers into clinical practice^28,29^.

To address this limitation, we have compiled a cohort of 290 patients procured from the Prostate Cancer Outcomes Cohort Study (ProCOC)^30^ to generate large-scale proteomic measurement of PCa tissue samples using data-independent acquisition mass spectrometry (DIA-MS). The data have been analysed through purpose-built computational workflows at the Australian Cancer Research Foundation International Centre for the Proteome of Human Cancer (ProCan^®^) in Westmead, Australia^31,32,33,34,35,36,37^. We have identified differentially expressed proteins and pathways involved in PCa development and biochemical recurrence (BCR), including the identification of possible new therapeutic targets. Further, we have built a protein-based signature, which showed better prognostic power than GG and was completely independent of it.

## RESULTS

### Proteomes of prostate tissue samples

A total of 290 PCa patients representing the full range of GG from GG1 to GG5 were selected from the ProCOC retrospective cohort^30^. Proteomes of 1,348 matched tumour and benign prostatic hyperplasia tissue sample runs from 278 patients were acquired and analysed with 12 patients being removed due to quality control (QC) steps. In each of the 31 batches, two controls (CTRL-A and CTRL-B) in duplicate were added to investigate technical variation, control quality, and assess reproducibility (**Fig. 1A; Methods; Supplementary Table 1; Supplementary Fig. 1)**. In this cohort, 198 of 278 patients had BCR data with a median follow-up of 59 months (**Supplementary Table 1**). Overall, most patients belong to GG2 (n = 135), followed by GG3 (n = 70; **Fig. 1A**). Although there was significant difference in outcome for GGs (*p*-value = 0.002), no significant difference was observed between GG2 and GG3, and GG4 unexpectedly showed the worst prognosis compared to all other GGs (**Supplementary Fig. 2**), reflecting the limitations of the GG system.

**Figure 1.**
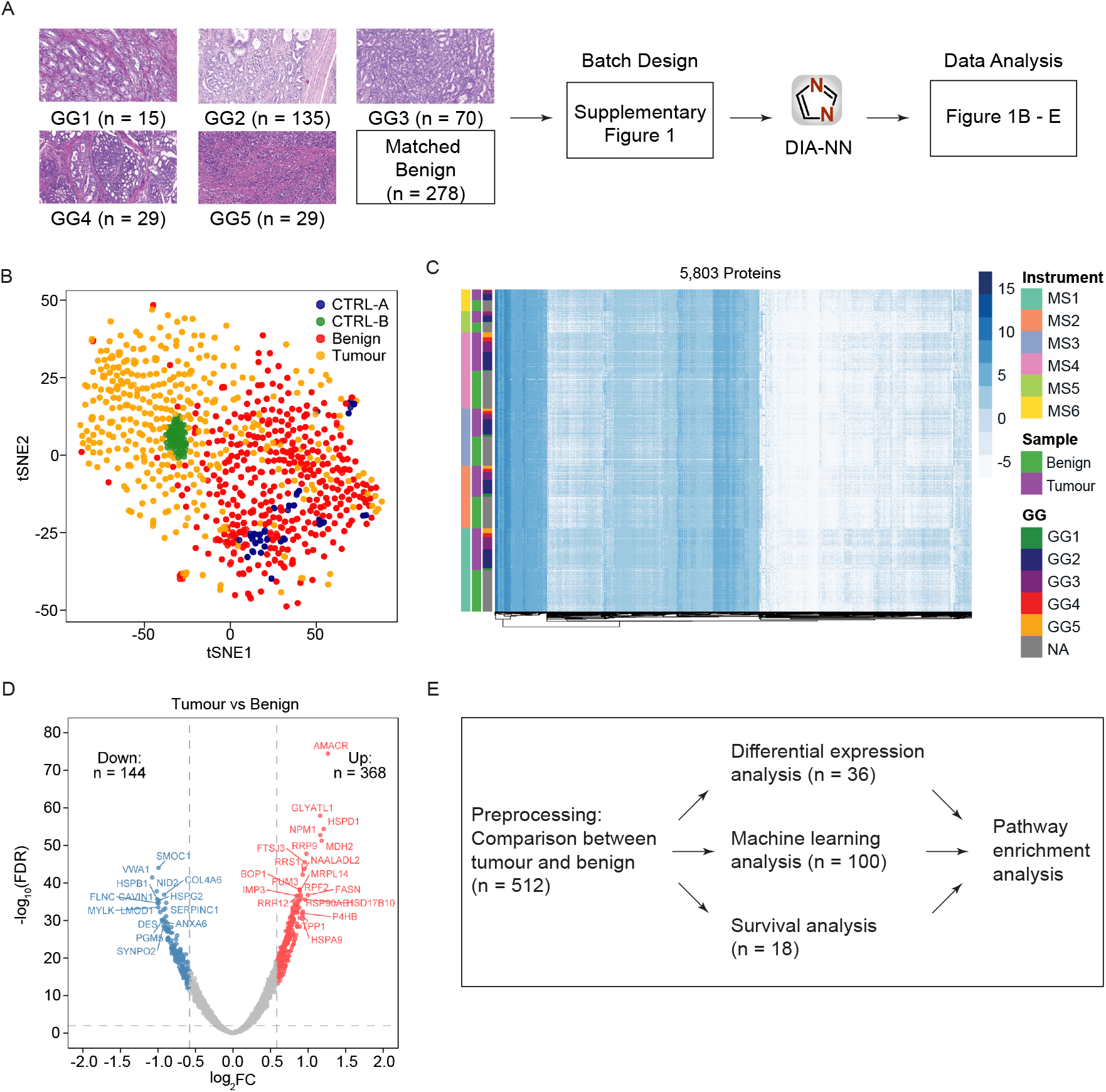
Proteomic analysis of PCa samples. **A.** An overview of the study design. Dataset consists of prostatic tumour and matched benign tissue samples from 278 patients. Proteomics data were collected for 277 tumour samples and 278 benign samples in duplicate from 278 patients. A total of 1,475 MS runs were analysed in 31 batches, including tumour, benign, CTRL-A and CTRL-B samples. The raw proteomics data were analysed by DIA-NN, quantifying 5,803 proteins. **B.** tSNE projection of protein data superimposed with colour annotation of sample types. **C.** Heatmap representation of the protein matrix with samples shown on the y-axis and proteins shown on the x-axis. The protein intensities were sorted first by the mass spectrometers, followed by tissue types and GGs. MS1 - MS6 indicate the six mass spectrometers. **D.** A volcano plot showing the upregulated (n = 368) and downregulated (n = 144) proteins in tumours with fold change (FC) > 1.5 and < 0.67 and Benjamini-Hochberg (BH) adjusted *p*-value < 0.01. Significant proteins are presented in red and blue colour, whereas other proteins are coloured in grey. **E.** Analysis pipeline employed in this study and the number of proteins identified in each analysis. A total of 512 tumour-enriched proteins were identified from the comparison between tumour and benign samples, followed by stratification of GG2 and GG3 using differential expression analysis and machine learning, and identification of a prognostic signature using survival analysis. Finally, pathway enrichment analyses were conducted for the significant sets of proteins.

Proteomic profiles of all samples including controls were acquired by DIA-MS in technical duplicate at ProCan^36^ using operating conditions that enable reproducible and high throughput data acquisition across six SCIEX™TripleTOF^®^6600 mass spectrometers^31,35^. We quantified 5,803 proteins (**Supplementary Fig. 3A**), with tumour samples showing a higher number of quantified proteins (average proteins per sample = 3,922) compared to benign samples (average proteins per sample = 3,587) (**Supplementary Fig. 3A**). The technical reproducibility of the cohort was evaluated by the Pearson correlation coefficient (Pearson’s r) among the sample replicates. There was a high degree of correlation between technical replicates of all samples with an average Pearson’s r of 0.94 (**Supplementary Fig. 3B**). Of the 5,803 proteins identified, >2,200 proteins were quantified in >90% of the samples and around 800 proteins were quantified in <20% of the samples (**Supplementary Fig. 3C**).

The t-distribution stochastic neighbour embedding (tSNE) analysis did not show batch effects from sample preparation. However, batch effects from different mass spectrometers appeared non-trivial without batch correction (**Supplementary Fig. 3D and 3E**). The tSNE analysis also showed a clear difference between benign and tumour samples (**Fig. 1B**). As expected, CTRL-A and CTRL-B samples are distinct from one another (**Fig. 1B**), implicating variation from both the mass spectrometer and sample preparation (CTRL-A) and variation from the mass spectrometer alone (CTRL-B). Tumour samples of high GGs (GG4 and GG5) were only partially separated from other groups, and separation of intermediate groups (GG2 and GG3) was barely visible (**Supplementary Fig. 3F**). A heatmap of the protein matrix showed distinct expression patterns of tumour and benign samples, however, no patterns were observed for GGs, which indicates that GG system alone does not explain the proteomic heterogeneity (**Fig. 1C**). Tumour and benign samples were compared by differential expression analysis as a data pre-processing step (**Fig. 1D**), resulting in the identification of 512 tumour-enriched proteins. These proteins were employed for the subsequent differential expression analysis, machine learning and survival analysis (**Fig. 1E**).

### Pre-processing by differential expression analysis between tumour and benign samples

To build a protein-based prognostic signature that stratifies GG2 and GG3 patients, we selected tumour-enriched proteins by performing a differential expression analysis between tumour and benign samples. In this pre-processing step, all tumour and matched benign samples were used with the full set of 5,803 proteins. The analysis resulted in 512 tumour-enriched proteins, of which 368 were upregulated and 144 were downregulated in tumour samples (**Fig. 1D**). The expression pattern of these differentially expressed proteins are shown in **Supplementary Fig. 4A** where proteins in the top cluster (upregulated proteins) exhibited considerably higher expression in tumour samples compared to benign samples (**Supplementary Fig. 4A**). Proteins in the bottom cluster were downregulated in tumour samples (**Supplementary Fig. 4A**).

Pathway enrichment analysis and protein-protein interaction (PPI) networks^38^ revealed that most of the upregulated pathways were related to ribosomal RNA processing, mitochondrial transmembrane transport, and protein folding (**Supplementary Fig. 4B and 4C**). When searched within the Hallmark gene sets^38^, the upregulated proteins were also found to be enriched in the MYC (proto-oncogene) targets V1 and V2 gene sets, which are known to be associated with tumour aggressiveness (**Supplementary Fig. 4D**).

Pathways and Gene Ontology (GO) processes that were significantly enriched in benign samples compared with tumour samples included muscle structure development, supramolecular fibre organization and response to elevated platelet cytosolic Ca2+ (**Supplementary Fig. 4B, 4C and 4D**). Of the top 20 differentially expressed proteins identified in tumour samples, four (MDH2, FASN, EPCAM, HSD17B10) are targetable by FDA-approved drugs, whereas two (AMACR and GLYATL1) are potentially targetable ^39^ and are of potential interest for future research.

The protein complexes identified in the PPI network, using the Molecular Complex Detection (MCODE) method showed upregulation of a large set of ribosomal proteins (both large and small subunits) that promote the process of protein translation, upregulation of proteins actively involved in the RNA metabolic process (RRS1, RPF2, BRIX1, RSL1D1), ribosome biogenesis (FTSJ3, DDX56, NPM3, GNL3, SNU13) and protein folding (HSPD1, HSPA9, HSPA5, PUM3) (**Supplementary Fig. 4E**). This is consistent with previous work showing the overexpression of proteins associated with cell adhesion, mitochondrial and ribosomal biogenesis and translation in PCa tissue samples^40^. Thus, we identified a list of differentially expressed proteins within tumour tissues for use in the downstream analyses, and identified a number of potentially important proteins and pathways in PCa.

### Stratification of GG2 and GG3 patients

To characterise the PCa samples from GG2 and GG3, we performed a differential expression analysis between the two GGs using the 512 tumour-enriched proteins. Of these, 35 proteins were significantly enriched in GG2 and one protein was enriched in GG3 (FC > 1.5 and < 0.67, *p*-value < 0.05, **Fig. 2A**). The significantly differentially expressed proteins formed two clusters based on their expression in GG2 and GG3 samples (**Fig. 2B**). As the set of significantly up- and down-regulated proteins was small, no significantly enriched pathway between GG2 and GG3 was identified. However, of the 35 upregulated proteins in GG2, two (TGFB1 and FLNA) are involved in androgen receptor pathways, three (FLNC, DES and LMOD1) have previously been associated with better prognosis in PCa^25,41,42,43,44,45^ four (PRKCA, ACTN1, AOC3 and LDHB) are targets for FDA-approved drugs^39^ and three (MYLK, FLNA, and FLNC) are potential drug targets^39^. The results suggested likely biological differences between GG2 and GG3 and identified several potential diagnostic and prognostic biomarkers that could be further investigated.

**Figure 2.**
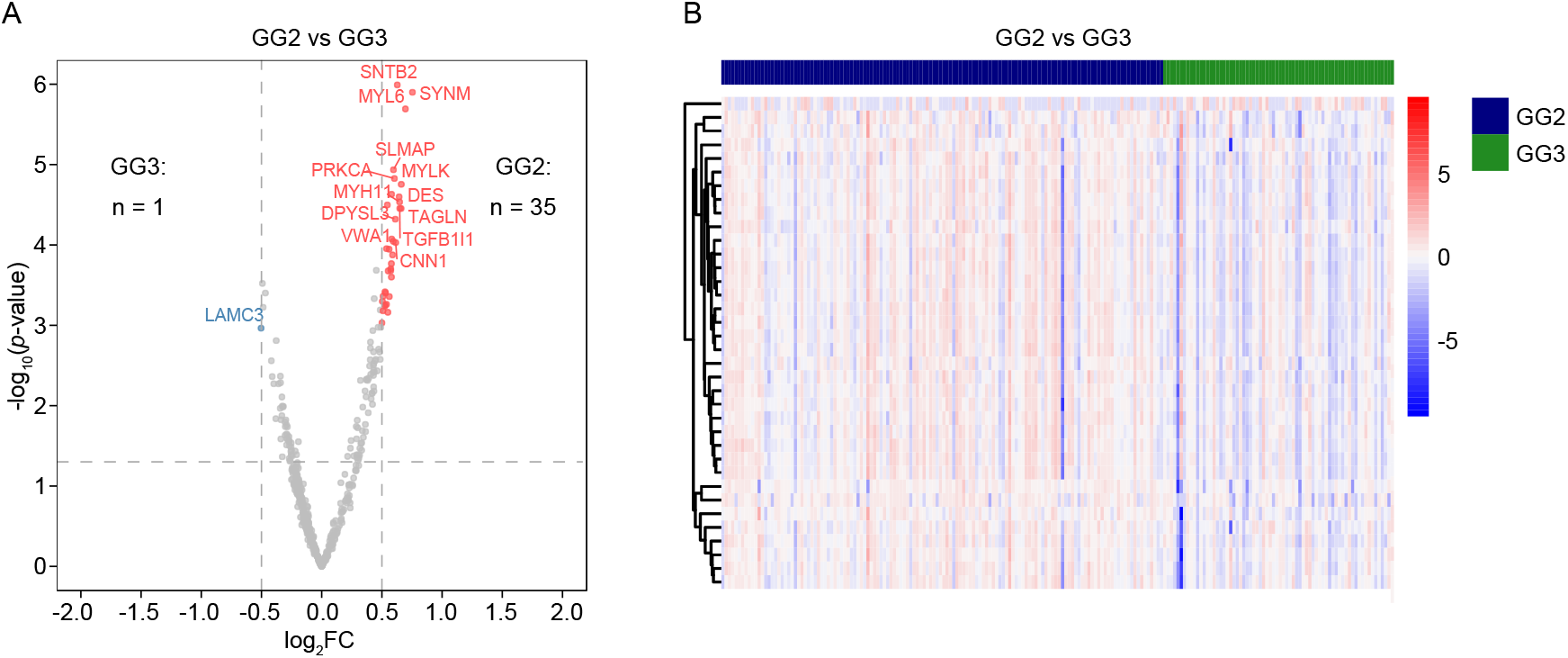
Differentially expressed proteins in GG2 vs GG3. **A.** A volcano plot showing the GG2 (n = 35) and GG3 (n = 1) enriched proteins in tumours with FC > 1.5 and < 0.67 and *p*-value < 0.01. Significant proteins are presented in red and blue colour, whereas other proteins are coloured in grey. Only a small number of proteins were found to be significant using differential expression analysis while most of them showed low FC. **B.** Heatmap representation of the expression levels of differentially expressed proteins between GG2 and GG3 samples shown in **A**. Expression data are converted to z-scores. Samples are shown on the x-axis whereas proteins are clustered on the y-axis.

To stratify GG2 against GG3 patients by machine learning, a dataset containing only GG2 and GG3 patients and the 512 tumour-enriched proteins was used. The results, aggregated over 1,000 Monte-Carlo cross-validation runs of an XGBoost classifier with 80% training and 20% testing splits, demonstrate that the difference between GG2 and GG3 can be predicted from protein intensities with high accuracy (**Fig. 3A**). The Receiver Operating Characteristics (ROC) curve of the best model had an Area Under the ROC (AUROC) of 0.89, with a mean AUROC of 0.74 (**Fig. 3A**). To obtain a reproducible list of top 20 most significant proteins in separating GG2 and GG3 samples, SHapley Additive exPlanations (SHAP) values were calculated over the entire cohort (**Fig. 3B**, see **Methods**).

**Figure 3.**
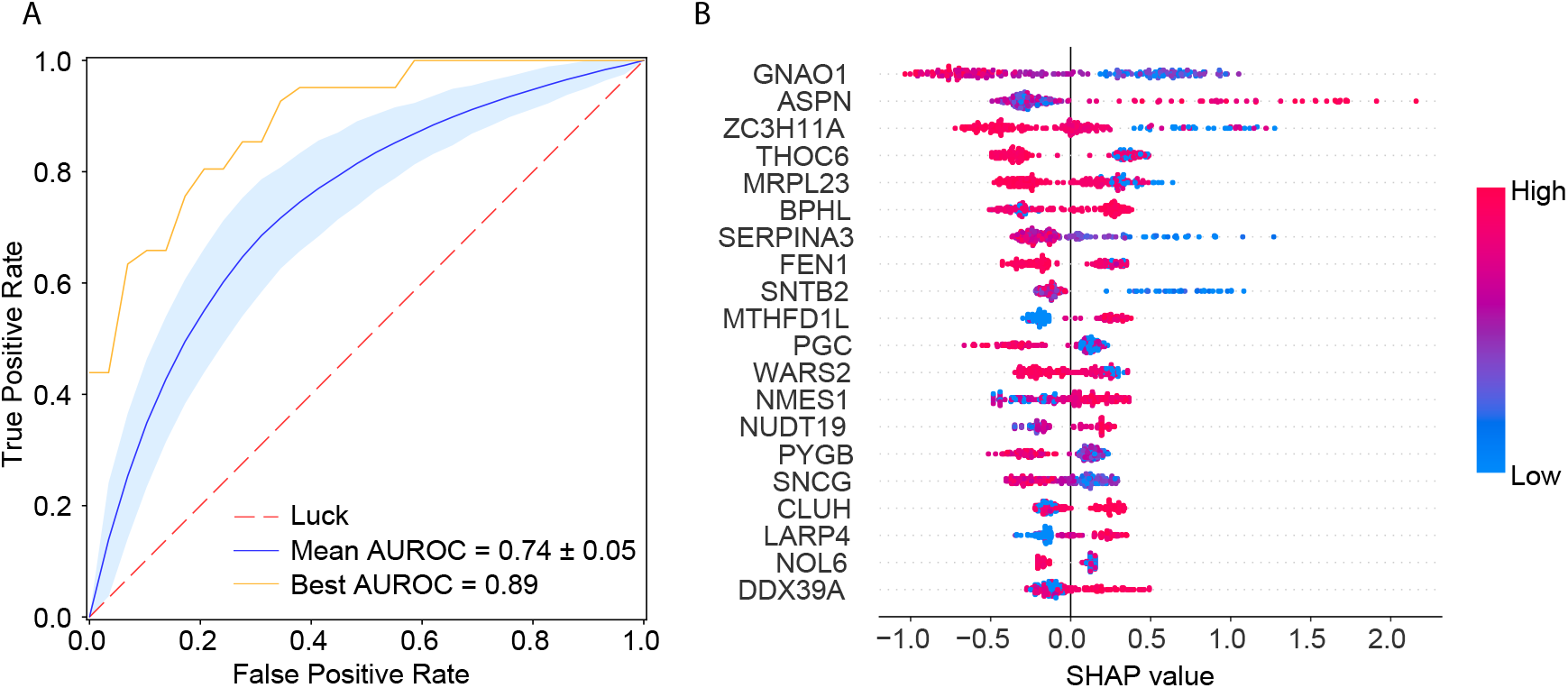
Machine learning of GG2 vs. GG3. **A.** ROC curves for the best and average models for predicting GG2 and GG3 samples based on 1000 Monte-Carlo runs by XGBoost. The red dashed line represents the random guess, the blue solid curve shows the mean ROC curve over 1000 Monte-Carlo runs, the blue band represents one standard deviation of the curves, and the orange curve shows the best ROC curve. **B.** SHAP values of the top 20 most significant proteins to distinguish between GG2 and GG3 samples, sorted (from top to bottom) by their respective absolute mean SHAP values. SHAP values of proteins in different samples are shown on the horizontal axis; the top 20 proteins are sorted (by importance) from top to bottom on the y-axis. The colours from blue to red indicate protein expression levels from low to high. The vertical zero-line (SHAP value = 0) is the line that has no impact on prediction, while the values on the left and right sides represent negative and positive impacts on prediction.

To study the dysregulated biological pathways in GG2 and GG3, a total of 127 proteins were selected by taking the union of 36 differentially expressed proteins **(Fig. 2A)** and the top 100 proteins from the machine learning that contains the top 20 proteins in **Fig. 3B**. Pathways enrichment analysis and PPI interactions from Reactome pathways^38^ for these proteins highlighted an overrepresentation of proteins involved in muscle structure, ECM organization and response to elevated platelet cytosolic Ca2+ pathways (**Fig. 4A**). When compared against the Hallmark gene sets, enrichment for proteins in the epithelial-mesenchymal transition gene sets was observed^38^ (**Fig. 4A**). The significant protein complexes identified in the PPI network using MCODE (**Fig. 4B**) included proteins involved in smooth muscle contraction (CALD1, TLN1, TPM2, TPM1, 4 myosin proteins), actin cytoskeleton proteins (ACTN4, MYO1C, FLNA), and mitochondrial translation (ribosomal subunit proteins). Most of these PPI proteins had high mean importance scores when GG2 samples were compared with GG3 samples. Our findings extend upon previous research showing that CALD1, TPM2 and TPM1 can be used as potential diagnostic biomarkers for PCa^46^. Although these findings are of biological interest, further modelling is required to better understand the biological pathways associated with each GG, and thus improve the GG prognostic performance.

**Figure 4.**
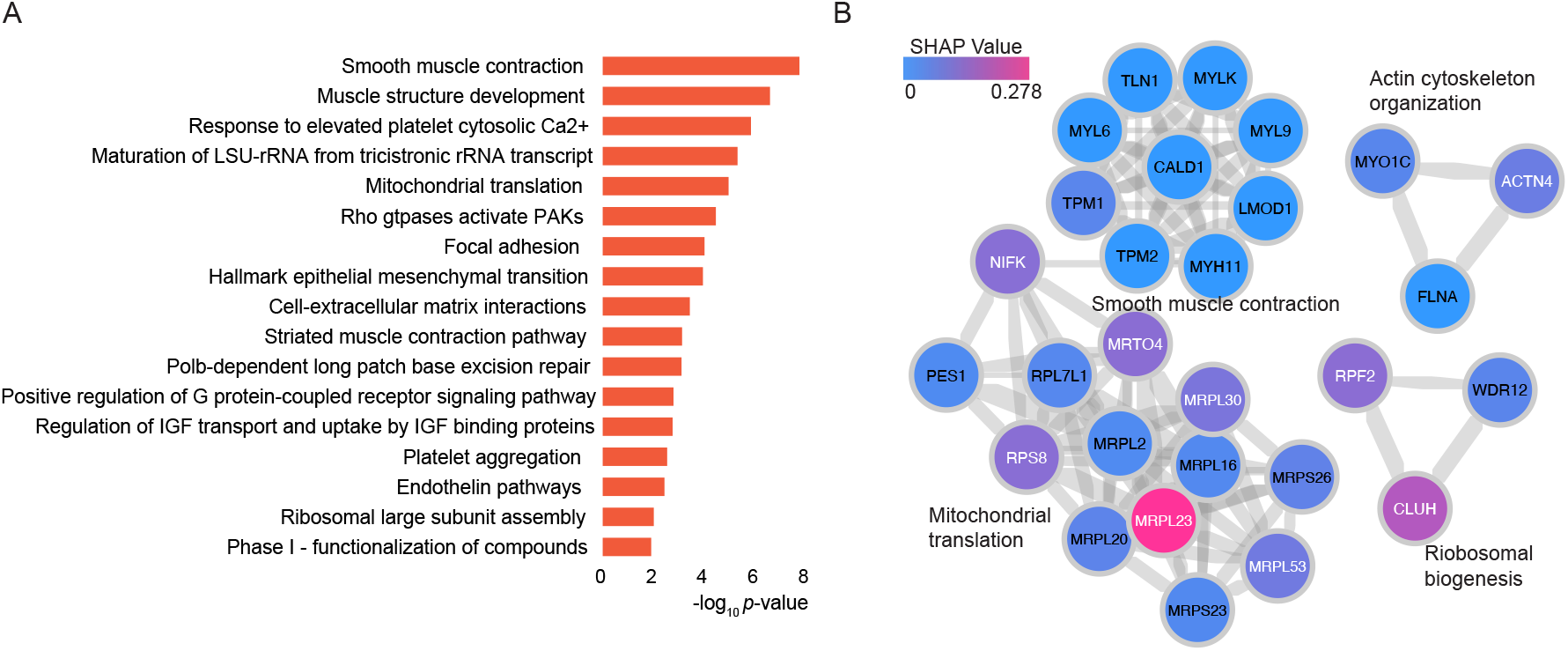
Differentially expressed proteins and pathways in GG2 and GG3 PCa. **A.** GO biological processes, Reactome pathways and hallmark gene sets enriched for the selected significant proteins. **B.** PPI network components obtained using MCODE algorithm, showing the enriched biological processes and proteins. Proteins are coloured according to the absolute mean SHAP values. The width of the edge (between nodes) indicates the strength of the connection. Functional description is provided beside each component.

### Protein-based prognostic signature for biochemical recurrence

To overcome one of the limitations of the GG system, exemplified in an inability to differentiate prognosis between GG2 and GG3 in our dataset (**Supplementary Fig. 2**), a protein-based signature was constructed. First, 100 runs of multivariate Cox regression^47^ with least absolute shrinkage and selection operator (LASSO) regularisation were performed on the 512 tumour-enriched proteins using 20-fold cross-validation (see **Methods**). For each run, a list of significant proteins was obtained, and a merged list of these proteins was collated and ranked according to the descending order of mean significance of individual proteins over all the 100 runs. A subset comprising the top 25 of these proteins was then used to model a multivariate Cox regression with recursive feature selection, yielding a final list of 18 proteins (**Fig. 5A**). Almost all of the 18 proteins were significantly associated with BCR with a concordance index (C-index)^48^ of 0.95 (**Fig. 5A**), indicating robust prognostic power from these proteins.

**Figure 5.**
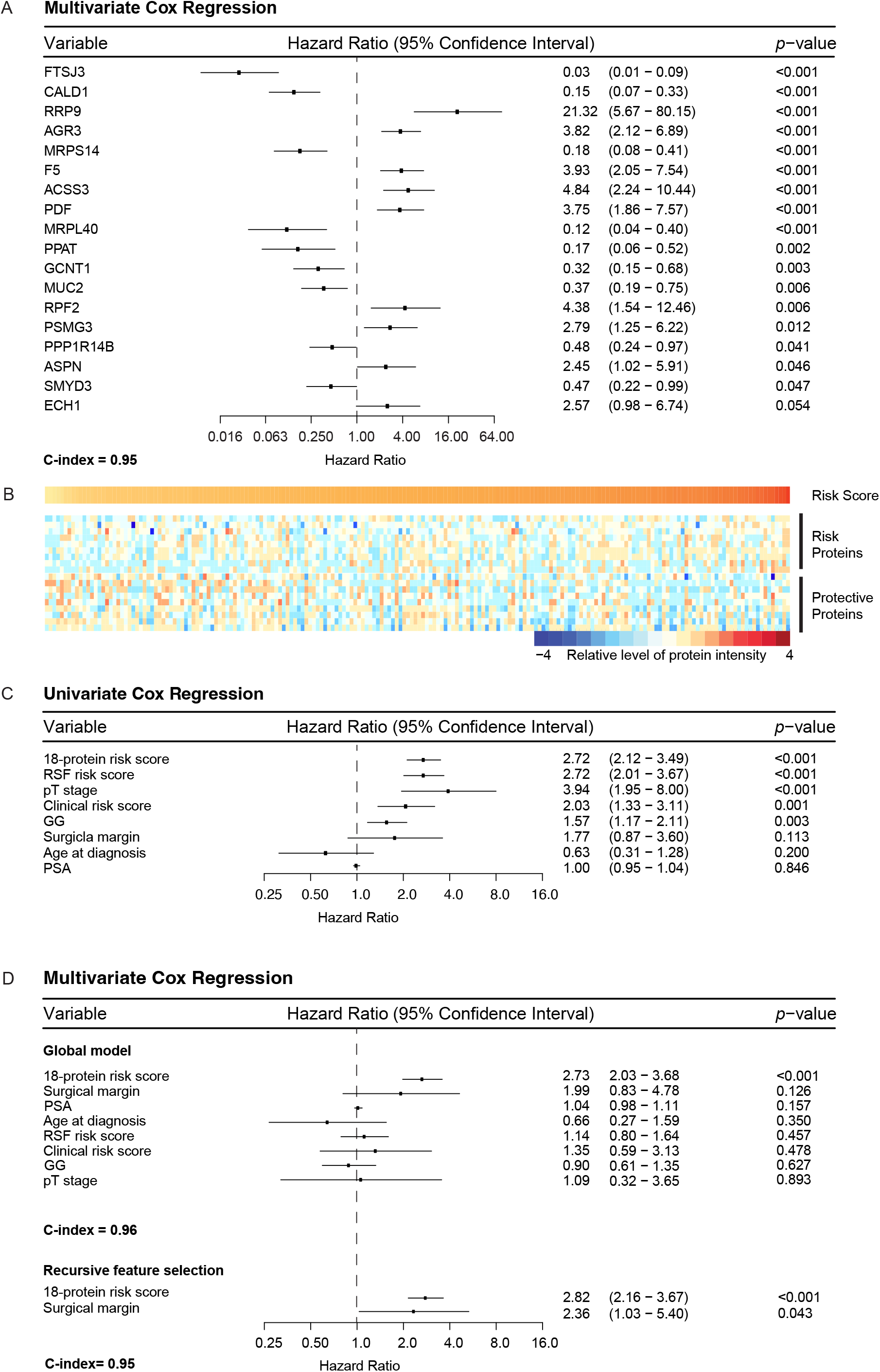
Survival Analysis of BCR-free survival (BCRFS) of PCa. **A.** Forest plot showing the 18 proteins with their individual hazard ratios, *p*-values, 95% CIs, and C-index of the final multivariate Cox model. **B.** Protein intensities for each PCa sample (according to the colour scale shown). The magnitude of the corresponding risk scores is represented by the scale bar. PCa samples with high risk scores expressed risk proteins, whereas samples with low risk scores expressed protective proteins. **C.** Forest plot comparing the importance of the 18-protein signature with RSF-based risk score and with other clinical variables using univariate Cox models. pT stage (pT1 vs pT2). Surgical margin (positive vs negative). Age at diagnosis (< 64 vs ≥ 64). **D.** Forest plot showing a simple multivariate Cox model that includes the 18-protein signature, RSF-based risk score and other clinical variables. With recursive feature selection, the 18-protein signature remains the most important variable, with a stable C-index (from 0.96 to 0.95).

A patient’s risk score was calculated as the sum of the intensities of each of the 18 proteins, multiplied by the corresponding regression coefficients (**Fig. 5A** and **Methods**). The midpoint of risk scores was used as the threshold to dichotomise patients to either a high-risk or a low-risk group. This two-step process gave rise to an 18-protein signature. To assess the prognostic power, the 18-protein signature was benchmarked with another signature calculated from the top 20 proteins identified by a random survival forests (RSF)^49^ model as well as with other clinicopathologic variables including GG, clinical risk, PSA, surgical margin, age at diagnosis, and pathological T stage (pT stage). The 18-protein signature showed the strongest association with BCR among all variables in the univariate Cox regression analysis (**Fig 5C**). This was also true in a multivariate Cox regression analysis after adjusting for the clinicopathologic variables and the 20-protein RSF signature independent of recursive feature selection (**Fig 5D**). This confirms that the 18-protein signature is not confounded by other clinicopathologic variables and can be considered an independent prognostic factor. The stable concordance index of all these models further suggests that the 18-protein signature can explain most of the association with BCR.

Moreover, our 18-protein signature was compared with the 20-protein RSF signature using a time-dependent ROC analysis, which measures how well an independent variable can differentiate between target classes at different time points in the study. The comparison of time-dependent ROC curves after 60 months for both risk scores showed an AUROC of 0.95 for the 18-protein signature and an AUROC of 0.82 for the RSF signature (**Supplementary Fig. 5A**). Further comparison demonstrated the higher predictive power of the 18-protein signature over time compared with the RSF signature (**Supplementary Fig. 5B**). RSF uses bootstrapped samples in each tree to avoid overfitting and generalises well on unseen test datasets^49^. For this reason, it is noteworthy that our 18-protein signature outperformed the RSF signature even in the absence of a validation dataset.

The dichotomised Kaplan-Meier curve with the low *p*-value (< 0.0001) indicated substantial predictive power by the 18-protein signature (**Fig. 6A**). Overall, there were more patients with GG2 and GG3 in our cohort compared with GG1, GG4 and GG5. Interestingly, the number of patients with GG2 and GG3 was equally distributed between the low-risk and high-risk groups (GG2: 55 and 50; GG3: 22 and 24, respectively), indicating that our protein-based signature is independent of GG. To confirm this, we applied the 18-protein signature within the group of patients including both GG2 and GG3 **(Fig. 6B)**, with GG2 only (**Fig. 6C**) and GG3 only (**Fig. 6D**). The 18-protein signature was able to identify a sub-group of patients with a higher risk of developing BCR within each GG, confirming its independence of GG and suggesting potential clinical utility.

**Figure 6.**
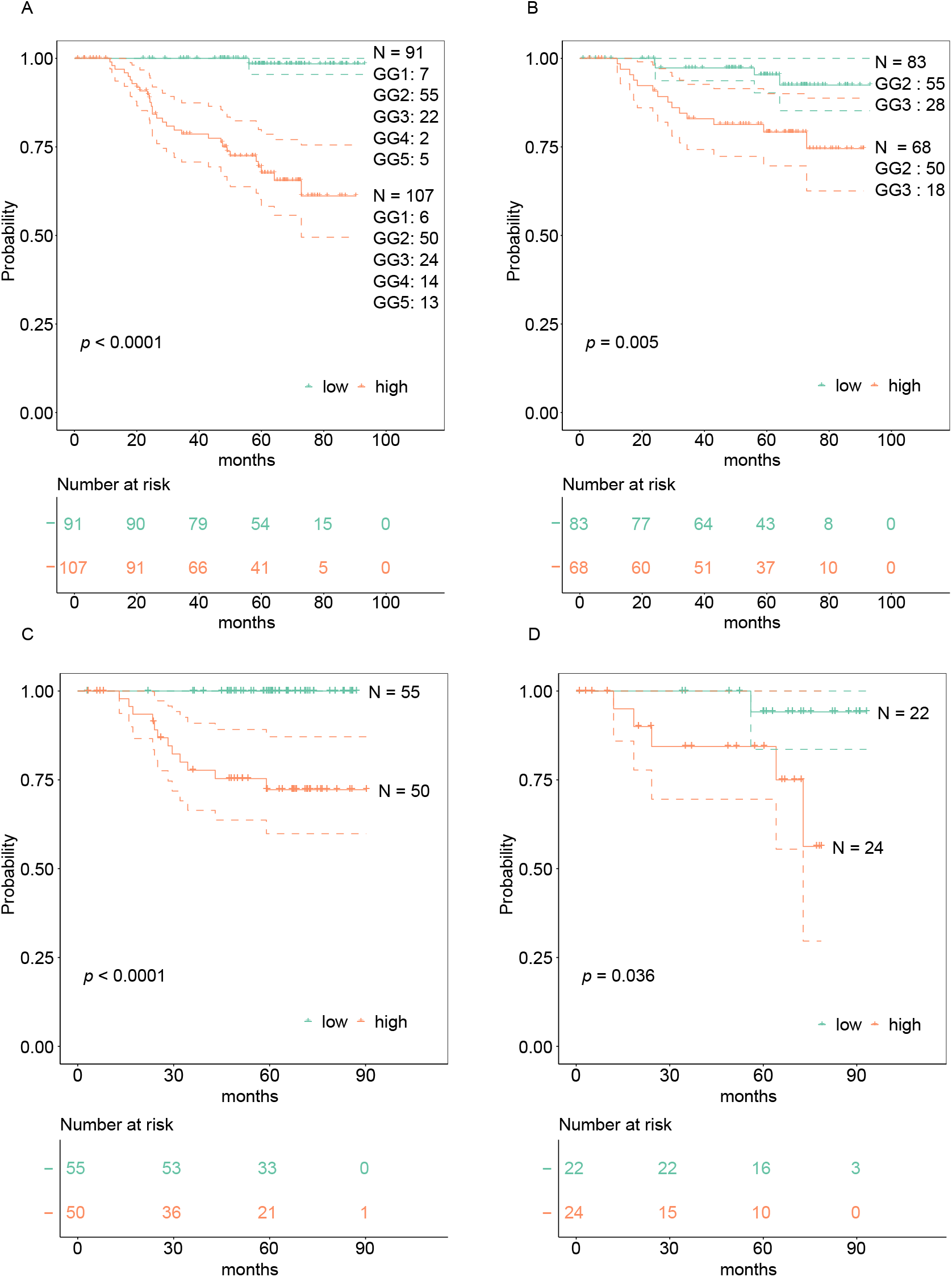
Kaplan-Meier (KM) curves for BCRFS. KM curves with 95% CIs of the low- and high-risk groups based on the 18-protein risk score, along with respective numbers of samples corresponding to each GG. Vertical lines illustrate patients who were censored at the time of their last clinical followup visit. The *p*-value shows significance of the difference between survival estimates evaluated by the log-rank test. Coloured values represent the number of patients in each group under risk. **A.** KM curves for PCa patients in all GGs. **B.** KM curves for PCa patients in GG2 and GG3. **C.** KM curves for PCa patients in GG2 only. **D.** KM curves for PCa patients in GG3 only.

By taking the union of the 18 signature proteins **(Fig. 5A)** and 26 proteins that were significantly associated with BCR in a univariate Cox regression model (*p*-value < 0.05), a total of 39 unique proteins (**Supplementary Fig. 6**) were analysed to study the association between biological pathways and BCR. Among these 39 proteins, five (F5, CALD1, RRP9, MUC2 and AGR3) were identified in common (see **Methods),** and six were related to androgen-regulated genes (F5, CALD1, TPM1, PUM3, ANXA4 and MYLK)^50^. Most of the 18 signature proteins were not involved in common biological pathways and thus contribute unique biological information. However, when including all 39 proteins, several enriched pathways were identified. This included muscle structure development (CALD1, MYL9, MYLK, TPM1) and rRNA metabolic processes (RRP9, PUM3, EARS2, RPF2, FTSJ3) (**Fig. 7A and 7B**). Of the total 39 proteins, two (F5 and ANXA4)^39^ are targetable by FDA-approved drugs and three (TMEM126B, EARS2, and MYLK) are potentially targetable^39^. Among the list of 26 proteins associated with BCR in the univariate Cox regression modelling, F5 (HR 1.7, 95% CI [1.2, 2.4]), TMEM126B (HR 1.5, 95% CI [1.1, 2.0]), and EARS2 (HR 1.9, 95% CI [1.1, 3.2]) were associated with increased risk of BCR, suggesting potential utility for further investigation as drug targets in clinical practice.

**Figure 7.**
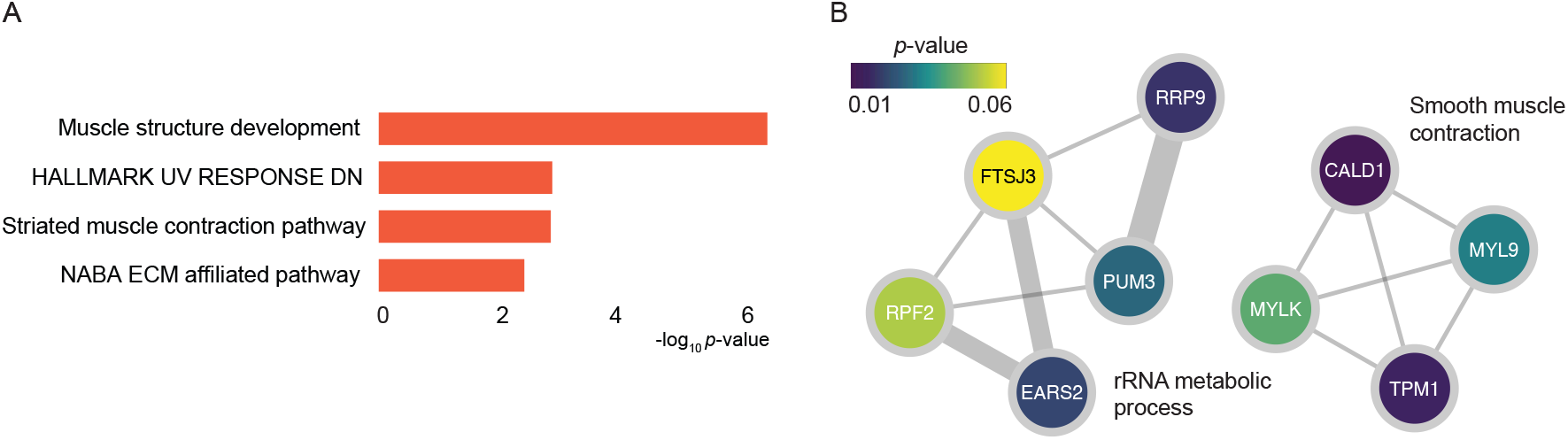
Significant biological pathways identified from univariate Cox regression model. **A.** GO biological processes, Reactome pathways and Hallmark gene sets enriched for the selected significant proteins. **B.** PPI network components obtained using the MCODE algorithm, showing the enriched biological processes and proteins. Proteins are coloured according to the *p*-values from the BCRFS analysis. The width of the edge (between nodes) indicates the strength of the connection. A functional description is provided next to each component.

## DISCUSSION

We performed a large-scale quantitative proteomic analysis from 278 PCa patients with primary tumour and matched benign tissue samples, each analysed in technical duplicate. We identified differentially expressed proteins and multiple signalling pathways related to PCa development and progression. In addition, we built an 18-protein signature that overcomes the limitations of GG in distinguishing between intermediate-risk PCa patients, and which has a higher prognostic value compared with the standard classification. We were also able to identify potential therapeutic targets that can be explored for their utility in the treatment of PCa. The main finding of this study is that patients with GG2 adenocarcinomas of the prostate (clinically the most common subgroup) could be significantly and independently divided into two subgroups with differential risk of BCR by our proteomic-based survival analysis, albeit an exploratory investigation.

The pathway enrichment analyses on tumour-enriched proteins showed that pathways related to protein folding, rRNA processing, ECM organisation, mitochondrial translation initiation, PCa development. Among the top 20 differentially expressed proteins, several proteins (AMACR, MDH2, FASN, HSD17B10) were involved in metabolic-related pathways^51,52,53^. Although few proteins were related to androgen (HSD17B10, F5, PUM3)^54^ and DNA damage repair (NPM1, FEN1)^54^ pathways, 16% of our 512 differentially expressed proteins overlapped with the over-expressed genes in PCa^55^. In addition, AMACR, FASN, IGFBP2, and PHB identified in our analysis are among biomarkers previously suggested for PCa diagnosis^40^.

Four of the top 20 differentially expressed proteins (MDH2, FASN, EPCAM, HSD17B10) are targetable with FDA-approved drugs, while two are potentially targetable proteins (AMACR and GLYATL1)^39^. AMACR was the top significantly upregulated protein in the tumour tissue. AMACR has a major role in fatty acid oxidation and has previously been found to be overexpressed in PCa at the proteomic and transcriptomic levels, confirming its validity as a potential biomarker^56,57,58,59^. Among the four FDA-approved targetable proteins, MDH2 is known to be overexpressed in PCa and castrate-resistant PCa (CRPC), highlighting its role in PCa progression^23^ and resistance to chemotherapy^60^. FASN is a key enzyme in de novo fatty acid synthesis and has been found to be overexpressed in CRPC and many other types of solid tumours ^51^. It is also associated with PCa progression, mainly through the activation of the PI3K/Akt/mTORC1 pathway, with a recent study suggesting the potential therapeutic benefit of its inhibition to overcome resistance to anti-androgen treatment^52^. EPCAM is a marker for cancer stem cells that are associated with cancer proliferation, adhesion and differentiation, and it is overexpressed in different types of cancer, including PCa^61^. In a meta-analysis, EPCAM overexpression was associated with a higher risk of BCR and the development of bone metastasis^62^. Finally, HSD17B10 is involved in different metabolic pathways, has an important role in regulating tissue androgen levels and may be involved in PCa progression through androgen-independent pathways^53^. Further studies will be required to confirm the value of these potential therapeutic targets in PCa management.

Our analyses identified 39 proteins significantly associated with BCR, of which five were listed in the Human Protein Atlas database^63^ either as FDA-approved targetable proteins (F5 and ANXA4) or potentially targetable proteins (TMEM126B, EARS2, and MYLK)^39^. None of these proteins overlapped with a published list of potential biomarkers for PCa aggressiveness or treatment resistance^56^. This may be due to the nature of our study cohort being a treatment-naïve patient population that was not yet exposed to anti-androgen treatment. However, three proteins (HNRNPA2B1, MRPS22, and PUM3) from our analysis were identified within the The Cancer Genome Atlas (TCGA) list of genes associated with poor prognosis^64^. Our results suggest the potential usefulness of F5, TMEM126B and EARS2 as potential therapeutic targets. Using PPI network analysis and tissue-specific gene co-expression network analysis, F5 was identified as one of the core genes in PCa^65^. Interestingly, F5 was also associated with an increased risk of breast cancer and the activation of the immune microenvironment^66^. TMEM126B is a complex I assembly factor that is critical for oxidative stress and inflammatory response^67^. Previous studies have demonstrated its role in response to chronic hypoxia through HIF-1-dependent mechanisms^68^. Although the role of TMEM126B in PCa is not fully explored, its interaction with HIF-1-dependent pathways, which play a critical role in PCa progression^69, 70^, warrants further exploration. EARS2 is involved in mitochondrial protein synthesis and was found to be associated with breast, pancreatic, renal and colorectal cancers^71,72^. There is some evidence of the co-expression of EARS2 with PALB2 in breast and pancreatic cancer and the association of their overexpression with poorer outcomes^72^. This finding suggests that PALB2 may also be involved in PCa progression and response to treatment^73,74,75^.

Despite the established prognostic value of the GG system and its use in PCa management, its limitations are well-recognised^8,9,12^. Previous studies have illustrated the differences between GG2 and GG3 on the metabolomic level, with higher intensity of phosphatidylcholines, and cardiolipins, among others, within GG3 samples, suggesting the involvement of differential biological pathway^17^. Similarly, Kawahara *et al.* performed proteomic analysis on 50 PCa tissue samples and identified a panel of 11 proteins that were associated with high-grade (GG4 and GG5) versus low-grade (GG1 and GG2) PCa^25^. Interestingly, this 11-protein panel was not able to distinguish samples within GG3^25^. In another study, a five-gene signature was constructed using data from the GEO and TCGA datasets, which was independent of the Gleason score when dichotomised as less or more than 7^76^. However, the prognostic power of this signature was not explored within each GG (especially the intermediate groups, GG2 and GG3).

In our analysis, there was an overlap between GG2 and GG3 in terms of their risk of developing BCR (**Supplementary Fig. 2**), reflecting the limitations of GG stratification. Our study identified 35 upregulated proteins in GG2 compared with GG3. These proteins were related to muscle structure development, epithelial-to-mesenchymal transition, metabolic pathways, and ECM interaction. As expected, most upregulated proteins are related to cancer genes^39^, with seven of them known to be enhanced in PCa (SYNM, DES, MYH11, TAGLN, CNN1, LMOD1, and PGM5)^39^. Four upregulated proteins within the GG2 group are FDA-approved drug targets (PRKCA, ACTN1, AOC3 and LDHB)^39^, and three are potential drug targets (MYLK, FLNA, and FLNC)^39^. In addition, several proteins that were upregulated in GG2 can be used as potential prognostic biomarkers that need further investigation. Of these, FLNC, a potential drug target that is involved in cell-extracellular matrix interaction has been associated with progression-free survival and lower risk of BCR^41,42^. DES, a cancer-enhanced gene that is involved in Aurora B signalling and striated muscle contraction, has been found to be underexpressed in PCa, and is associated with better prognosis^43,44,45^. Finally, LMOD1, a PCa-enhanced gene has lower expression in high-grade and metastatic PCa^25^. Further research is required to determine the utility of those proteins as prognostic biomarkers at the time of PCa diagnosis.

To overcome the limitations of GG, we have built a protein-based signature and explored its prognostic power together with and in comparison with GG. Our 18-protein signature identified patients at higher risk of developing BCR with high accuracy. Its prognosis was maintained even after adjusting for other clinical variables, including GG, pT-stage, and baseline PSA. In addition, the 18-protein signature was independent of GG, being able to identify patients at a higher risk of developing BCR within each of the GG2 and GG3 groups separately. This distinction is of considerable clinical importance, considering the recent BCR management guidelines, which depend only on GG and PSA doubling time^77^. Further exploration of this protein-based signature for patients planned for active surveillance would be useful considering its potential ability to identify patients at higher risk of progression independent of their clinical risk score (PSA, GG and pT-stage)^3^. Our results both complement and extend upon recent proteomic studies in PCa^27^. The novel contribution of our work is first in presenting a substantially larger cohort size (n = 278) than previous studies, which typically comprise <100 patients^27^. Second, our study is able to identify potential novel therapeutic targets and build a prognostic signature that is completely independent of the GG, with the ability to identify patients at higher risk of developing BCR within the relatively indolent GG2. Although BCR is a problematic endpoint^78^, evidence suggests that patients who develop BCR are at higher risk of developing clinical progression^79^. It will be important to further investigate and validate the utility of our 18-protein signature on selecting the group of patients at higher risk of clinical progression and poorer survival. Finally, our dataset will serve as an important public resource for the scientific community seeking to understand the proteomic landscape in PCa.

This study is hampered by the unavailability of metastatic relapse and mortality data and the smaller number of patients within the GG1, GG4 and GG5 groups, which prevented us from confirming the prognostic value of the 18-protein signature within these GG groups. Although we did not have access to a validation cohort to verify our findings at the time of these analyses, our data will become an important resource for any future work requiring a validation data set. Showing that our 18-protein signature had higher significance and AUROC as compared with the 20-proteins RFS signature does provide a level of confirmation because the RSF model works on selecting bootstrapped samples in each tree while computing the importance of proteins. This process mimics internal cross-validation, avoids overfitting, and has been shown to generalise well on future data^49^.

We conclude that PCa proteomic analysis is a promising tool for understanding the biological pathways associated with PCa development and progression. Our analysis has identified several novel therapeutic targets, and possible diagnostic and prognostic biomarkers that can be further investigated in pre-clinical and clinical studies. Importantly, we have also built an 18-protein signature that was predictive of BCR and is independent of GG. Further work is required to first validate our findings in an independent cohort and then to integrate them into clinical practice.

## ONLINE METHODS

### Biospecimen collection and pathology and clinical data

The sample collection of this study was approved by the Cantonal Ethics Committee of Zürich (KEK-ZH-No. 2008-0040). Detailed information on the patients and samples has been published^15^ (PMID: 33317623) and is also provided in Table S1. Tumour tissue samples were fixed with formalin and embedded with paraffin.

### Sample preparation and mass spectrometric acquisition

About 0.5 mg of FFPE tissue was punched from the sample, weighed and processed for each biological replicate via the workflow as described previously^34^.

An Eksigent nanoLC 425 HPLC operating in microflow mode, coupled online to a 6600 Triple TOF (SCIEX) was used for the analyses. The peptide digests (2 μg) were injected onto a C18 trap column (SGE TRAPCOL C18 G 300 μm x 100 mm) and desalted for 5 min at 8 μL/min with solvent A (0.1% [v/v] formic acid). The trap column was switched in-line with a reversed-phase capillary column (SGE C18 G 250 mm × 300 μm ID 3 μm 200 Å), maintained at a temperature of 40°C. The flow rate was 5 μL/min. The gradient started at 2 % solvent B (99.9% [v/v] acetonitrile, 0.1% [v/v] formic acid) and increased to 35% over 69 min. This was followed by an increase of solvent B to 95% over 4 min. The column was washed with 95% solvent B for 5 min, then decreased to 2% solvent B over 3 min followed by a 13 min column equilibration step with 98% solvent A. For SWATH acquisition peptide spectra were analysed using the Triple TOF 6600 system (SCIEX) equipped with a DuoSpray source and 50 μm internal diameter electrode and controlled by Analyst 1.7.1 software. The following parameters were used: 5500 V ion spray voltage; 25 nitrogen curtain gas; 100°C TEM, 20 source gas 1, 20 source gas 2 with 100 variable windows, as per SCIEX technical notes. The parameters were set as follows: lower m/z limit 350; upper m/z limit 1250; 150 ms acquisition time, window overlap (Da) 1.0; CES was set at 5 for the smaller windows, then 8 for larger windows; and 10 for the largest windows. MS2 spectra were collected in the range of m/z 100 to 2000 for 30 ms in high-resolution mode and the resulting total cycle time was 3.2s.

### Proteomic data analysis

We analysed 278 out of 290 PCa patients whose malignant tissue samples were classified by pathologists alongside matched benign tissue. A total of 12 patients were removed after QC. The entire cohort was then divided into 31 batches, with each containing between 15 and 29 samples including two control samples (CTRL-A, n = 62 and CTRL-B, n = 62) for QC and the evaluation of reproducibility (**Supplementary Fig. 1)**. The samples were analysed in technical duplicate in different mass spectrometers in ProCan^31,36^.

From each patient, a malignant tissue sample and its matched benign sample were processed using pressure cycling technology (PCT)^80^ in technical duplicates, and randomly selected samples were processed with both biological replicates and technical replicates. The samples were processed in 31 batches, each containing a reference sample (CTRL-A) of a homogeneous PCa tissue sample that could account for technical variation introduced during the entire PCT-SWATH-MS sample processing methodology, and a reference sample of a homogeneous prostate tissue digest (CTRL-B) that could account for technical variation introduced during SWATH-MS.

#### DIA-based spectral library generation

DIA-MS data in wiff file format were collected for 1,475 runs and were processed using DIA-NN (version 1.8)^81^. A spectral library was generated using 1,475 DIA-MS runs and consisted of 9230 proteins and 89,408 peptides. The spectral library was used to search the complete cohort of 1,475 runs.

#### Data extraction

DIA-NN was implemented using RT-dependent normalization and with parameters given below:

~~~
-report-lib-info --out step3-out.tsv --qvalue 0.01 --pg-level 1 --mass-acc-
ms1 40 --mass-acc 40 --window 9 --int-removal 1 --matrices --temp . --smart-
profiling --peak-center
~~~

Data were then filtered to retain only precursors from proteotypic peptides with Global.Q.Value ≤ 0.01. Proteins were then quantified using maxLFQ, with default parameters^82^ and implemented using the DIA-NN R Package (https://github.com/vdemichev/diann-rpackage). Data were then log2-transformed. There were 1,475 mass spectrometry runs with 669 benign and 679 tumour samples. For downstream analysis, a final protein matrix with only benign and tumour samples (n = 1,348 samples) was used. The protein matrix showed an average of 35% missingness per individual sample. Missing values in this dataset were then imputed with a constant lower than the minimum value of the whole protein matrix to maintain the distinction between missing values and protein intensities. Sample replicates were merged. The imputed protein matrix was z-score standardized and was then used as input for further analyses.

#### Batch effect analysis

The tSNE analysis of the data were performed on the final protein data matrix with 5,803 proteins. The instrument batch effect was observed as samples were run on six different mass spectrometers. The tSNE-based two-dimensional visualization of protein data showed that the instrument batch effect was corrected after the built-in normalization method in the software suite DIA-NN (**Supplementary Fig. 3D and 3E**).

### Differential proteomic analysis

Differential expression analysis between tumour and benign samples was performed on all 5,803 proteins, and analysis between GG2 and GG3 samples was performed on 512 tumour-enriched proteins. Empirical Bayes moderated t-statistics, packaged in the Limma R package version 3.54.1, was performed to compute the *p*-value of protein intensity between the two classes. Tumour-specific significantly expressed proteins were selected at Benjamini-Hochberg (BH) adjusted *p*-value < 0.01 and with log fold change (FC) (expressed as the difference in the group means) cut-off of ±0.5 (FC > 1.5 and < 0.67), whereas GG2 and GG3-specific differentially expressed proteins were selected at *p*-value < 0.01 and with log FC (expressed as difference in the group means) cut-off of ±0.5 (FC > 1.5 and < 0.67). Heatmaps were generated using the R package pheatmap version 1.0.12. The complete linkage clustering algorithm was used along with Euclidean distance as the distance measure.”

### Survival analysis

#### Finding a proteomic signature

The protein dataset, containing the 512 tumour-enriched proteins, was used as the input for the survival analysis. To reduce the number of important proteins, 100 runs of multivariate Cox regression with LASSO regularization were executed on the whole dataset. The LASSO regularization hyperparameter in each run was tuned using 20-fold cross-validation. Each run returned a list of proteins with non-zero coefficients. These lists were then combined into a list of unique proteins, which was then ranked according to the mean importance of the individual proteins (average absolute coefficient over 100 runs) in descending order. The top 25 of these proteins were then used in a multivariate Cox model with recursive feature selection, which yielded the final 18 proteins. These 18 proteins were then used to construct the proteomic risk score (*S_j_*), for the *j*_th_ patient, as below:

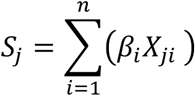

where *n* is total number of proteins; *β_i_* is the co-efficient of the *i*_th_ protein and *X_ji_* the intensity of the *i*_th_ protein, in the *j*_th_ patient.

#### Analysing performance of the proteomic risk score

The performance of the risk scores was analysed in multiple ways. First, patients were dichotomized into low and high-risk groups using midpoint of the range of risk scores as a threshold, and their Kaplan-Meier (KM) curves were then plotted. Differences between survival estimates were evaluated by the log-rank test and *p*-values were reported. The number of samples corresponding to each GG falling in both low- and high-risk groups were counted to analyse how well the KM curves justified categorization based on GG. Furthermore, to check its performance in GG2 and GG3 patients, KM curves for the dichotomized risk score were plotted in both combined as well as separate GG2 and GG3 patients.

The C-index is a measure of rank correlation between the predicted risk score and the observed time points. For instance, if the predicted risk score of a sample is higher than that of another, and the observed time point for that sample is earlier than that of the other sample, then the predictions and observations are said to be concordant.

### Functional enrichment analysis

Functional and pathways enrichment analysis of significantly expressed proteins was performed using Metascape^38^ along with the entire set of 5,803 proteins as the background gene set. The gene ontology (GO) biological processes, Reactome pathways and Hallmark gene sets enriched in dysregulated proteins were acquired. The input parameters were *p*-value < 0.05, minimum gene count of 3 and enrichment factor > 1. The *p*-values are calculated based on accumulative hypergeometric distribution and are adjusted using BH correction. For tumour versus benign, statistically significant enriched terms were selected at adjusted *p*-value (*q*-value or FDR) of 0.05 (−log10 FDR > 1.3), whereas for GG2 vs. GG3 comparisons, statistically significant enriched terms were selected at *p*-value of 0.05 (−log10 *p*-value > 1.3).

### PPI enrichment analysis

PPI enrichment analysis was performed using the Metascape^38^ by incorporating the data from STRING and BioGrid databases. As a result, a network of subsets of proteins is formed where proteins in the input list form physical interaction with at least one other member in the list. In order to identify the functional protein complexes for the differentially expressed proteins, the Molecular Complex Detection (MCODE) algorithm was applied within the Metascape^38^. MCODE detects and generates the significant protein complexes (*p*-value < 0.05) with minimum three proteins and maximum 500 proteins and provides the functional description for each complex. Using the MCODE algorithm, proteins and protein complexes that are enriched in the significantly dysregulated pathways were identified. The protein networks were visualized using Cytoscape^83^ where nodes represent the proteins and edges represent the connections between the nodes. The size of the node in a complex shows the MCODE score whereas width of the edge shows the strength of the connection.

### Machine learning

The protein dataset with 512 tumour-enriched proteins was used as the input in this analysis. Since the number of patients is not large, a single train and test split of the dataset will lead to biased conclusions. Therefore, we decided to draw our conclusion based on results aggregated from multiple Monte-Carlo runs of XGBoost classifier with random train and test splits. We used 1000 runs of Monte-Carlo cross-validation on a random XGBoost classifier with 300 base learners and the rest of the hyperparameters set to defaults (Python package “xgboost”). In each Monte-Carlo run, the dataset was split randomly into 80% training and 20% test sets, stratified by the target variable GG (GG2 vs. GG3). The test results from all the 1000 runs were then aggregated and the expected performance was reported.

## DATA AVAILABILITY

Detailed information will be provided in the peer-reviewed article.

## AUTHOR CONTRIBUTIONS

QZ, AB, PGH, RRR, PJR, RA, TG, and PJW contributed to the study design. QZ, DR, NJR, JHR, CP, TH, CF, and PJW contributed to sample procurement and clinical data organization. YZ, NL, PGH, and TG contributed to sample preparation and analysis. QZ, SR, ATA, ZN, AA, RCP, ML, TZ, BW, GBC, YW, XD, MRM, WS, MB, JFN, AB, and TG contributed to the data analysis. QZ, AA, ZN, AA, and PGH wrote the manuscript. All co-authors contributed to writing and approval of the final manuscript.

## ACKNOWLEDGEMENT

We thank Matteo Manica, Patrick Pedrioli, Goerge Rosenberger, Hannes Rost and Jelena Cuklina for helpful comments and discussions. ProCan is supported by the Australian Cancer Research Foundation, Cancer Institute New South Wales (NSW) (2017/TPG001, REG171150), NSW Ministry of Health (CMP-01), The University of Sydney, Cancer Council NSW (IG 18-01), Ian Potter Foundation, the Medical Research Future Fund (MRFF-PD), National Health and Medical Research Council (NHMRC) of Australia European Union grant (GNT1170739, a companion grant to support the European Commission’s Horizon 2020 Program, H2020-SC1-DTH-2018-1, ‘iPC - individualizedPaediatricCure’ [ref. 826121]), and National Breast Cancer Foundation (IIRS-18-164). The work at ProCan was done under the auspices of a Memorandum of Understanding between Children’s Medical Research Institute and the U.S. National Cancer Institute’s International Cancer Proteogenomics Consortium (ICPC), that encourages cooperation among institutions and nations in proteogenomic cancer research in which datasets are made available to the public. RCP and PJR are supported by the NHMRC Fellowships (GNT1138536 and GNT1137064, respectively). WS is supported by the National Natural Science Foundation of China (grant 62102248). MB is supported by the SNSF SystemsX.ch fellowship (TPdF 2013/135). MRM is supported by the European Union’s Horizon 2020 research and innovation programmes (668858 and 826121). RA is supported in part by SystemsX.ch the Swiss initiative for systems biology.

## REFERENCE

1. Sung H, et al. Global Cancer Statistics 2020: GLOBOCAN Estimates of Incidence and Mortality Worldwide for 36 Cancers in 185 Countries. CA Cancer J Clin 71, 209–249 (2021).

2. National Comprehensive Cancer Network (NCCN). Prostate Cancer Version 3.) (2022).

3. D’Amico AV, et al. Biochemical outcome after radical prostatectomy or external beam radiation therapy for patients with clinically localized prostate carcinoma in the prostate specific antigen era. Cancer 95, 281–286 (2002).

4. van Leenders GJ, Verhoef EI, Hollemans E. Prostate cancer growth patterns beyond the Gleason score: entering a new era of comprehensive tumour grading. Histopathology 77, 850–861 (2020).

5. Chan TY, Partin AW, Walsh PC, Epstein JI. Prognostic significance of Gleason score 3+4 versus Gleason score 4+3 tumor at radical prostatectomy. Urology 56, 823–827 (2000).

6. Epstein JI, Egevad L, Amin MB, Delahunt B, Srigley JR, Humphrey PA. The 2014 International Society of Urological Pathology (ISUP) Consensus Conference on Gleason Grading of Prostatic Carcinoma: Definition of Grading Patterns and Proposal for a New Grading System. Am J Surg Pathol 40, 244–252 (2016).

7. Mathieu R, et al. Prognostic value of the new grade groups in prostate cancer: a multi-institutional European validation study. Prostate cancer and prostatic diseases 20, 197–202 (2017).

8. Srigley JR, et al. Controversial issues in Gleason and International Society of Urological Pathology (ISUP) prostate cancer grading: proposed recommendations for international implementation. Pathology 51, 463–473 (2019).

9. Van Leenders GJ, et al. The 2019 International Society of Urological Pathology (ISUP) consensus conference on grading of prostatic carcinoma. The American journal of surgical pathology 44, e87 (2020).

10. Zhong Q, et al. Image-based computational quantification and visualization of genetic alterations and tumour heterogeneity. Sci Rep 6, 24146 (2016).

11. Zhong Q, et al. A curated collection of tissue microarray images and clinical outcome data of prostate cancer patients. Sci Data 4, 170014 (2017).

12. van der Slot MA, et al. Inter-observer variability of cribriform architecture and percent Gleason pattern 4 in prostate cancer: relation to clinical outcome. Virchows Archiv 478, 249–256 (2021).

13. Shao W, et al. Comparative analysis of mRNA and protein degradation in prostate tissues indicates high stability of proteins. Nat Commun 10, 2524 (2019).

14. Guo T, et al. Multi-region proteome analysis quantifies spatial heterogeneity of prostate tissue biomarkers. Life Sci Alliance 1, (2018).

15. Charmpi K, et al. Convergent network effects along the axis of gene expression during prostate cancer progression. Genome Biol 21, 302 (2020).

16. Zerhouni E, Prisacari B, Zhong Q, Wild P, Gabrani M. A computational framework for disease grading using protein signatures. In: 2016 IEEE 13th international symposium on Biomedical Imaging (ISBI)). IEEE (2016).

17. Randall EC, et al. Molecular characterization of prostate cancer with associated Gleason score using mass spectrometry imaging. Molecular Cancer Research 17, 1155–1165 (2019).

18. Peng Z, et al. Improving the prediction of prostate cancer overall survival by supplementing readily available clinical data with gene expression levels of IGFBP3 and F3 in formalin-fixed paraffin embedded core needle biopsy material. 11, e0145545 (2016).

19. Peng Z, et al. Operator dependent choice of prostate cancer biopsy has limited impact on a gene signature analysis for the highly expressed genes IGFBP3 and F3 in prostate cancer epithelial cells. 9, e109610 (2014).

20. Peng Z, et al. An expression signature at diagnosis to estimate prostate cancer patients’ overall survival. 17, 81–90 (2014).

21. Carneiro A, et al. Are localized prostate cancer biomarkers useful in the clinical practice? Tumor Biology 40, 1010428318799255 (2018).

22. Campistol M, Morote J, Regis L, Celma A, Planas J, Trilla E. Proclarix, A New Biomarker for the Diagnosis of Clinically Significant Prostate Cancer: A Systematic Review. Mol Diagn Ther 26, 273–281 (2022).

23. Latonen L, et al. Integrative proteomics in prostate cancer uncovers robustness against genomic and transcriptomic aberrations during disease progression. Nature communications 9, 1–13 (2018).

24. Garcia-Marques F, et al. Protein signatures to distinguish aggressive from indolent prostate cancer. Prostate, (2022).

25. Kawahara R, et al. Tissue Proteome Signatures Associated with Five Grades of Prostate Cancer and Benign Prostatic Hyperplasia. Proteomics 19, e1900174 (2019).

26. Zhu Y, Aebersold R, Mann M, Guo T. SnapShot: Clinical proteomics. Cell 184, 4840–4840.e4841 (2021).

27. Sadeesh N, Scaravilli M, Latonen L. Proteomic landscape of prostate cancer: The view provided by quantitative proteomics, integrative analyses, and protein interactomes. Cancers 13, 4829 (2021).

28. Xiao Q, et al. High-throughput proteomics and AI for cancer biomarker discovery. Adv Drug Deliv Rev 176, 113844 (2021).

29. Poulos RC, Cai Z, Robinson PJ, Reddel RR, Zhong Q. Opportunities for pharmacoproteomics in biomarker discovery. Proteomics, e2200031 (2022).

30. Umbehr M, et al. ProCOC: the prostate cancer outcomes cohort study. BMC Urol 8, 9 (2008).

31. Poulos RC, et al. Strategies to enable large-scale proteomics for reproducible research. Nat Commun 11, 3793 (2020).

32. Gonçalves E, et al. Pan-cancer proteomic map of 949 human cell lines. Cancer Cell 40, 835–849.e838 (2022).

33. Seneviratne AJ, et al. Improved identification and quantification of peptides in mass spectrometry data via chemical and random additive noise elimination (CRANE). Bioinformatics 37, 4719–4726 (2021).

34. Zhu Y, et al. High-throughput proteomic analysis of FFPE tissue samples facilitates tumor stratification. Mol Oncol 13, 2305–2328 (2019).

35. Manda SS, Noor Z, Hains PG, Zhong Q. PIONEER: Pipeline for Generating High-Quality Spectral Libraries for DIA-MS Data. Curr Protoc 1, e69 (2021).

36. Tully B, Balleine RL, Hains PG, Zhong Q, Reddel RR, Robinson PJ. Addressing the Challenges of High-Throughput Cancer Tissue Proteomics for Clinical Application: ProCan. Proteomics 19, e1900109 (2019).

37. Cadow J, Manica M, Mathis R, Guo T, Aebersold R, Rodriguez Martinez M. On the feasibility of deep learning applications using raw mass spectrometry data. Bioinformatics 37, i245–i253 (2021).

38. Zhou Y, et al. Metascape provides a biologist-oriented resource for the analysis of systems-level datasets. 10, 1–10 (2019).

39. Uhlén M, et al. Tissue-based map of the human proteome. Science 347, 1260419 (2015).

40. Iglesias-Gato D, et al. The proteome of primary prostate cancer. European urology 69, 942–952 (2016).

41. Pachynski RK, et al. Single-cell Spatial Proteomic Revelations on the Multiparametric MRI Heterogeneity of Clinically Significant Prostate CancerIntegrated Proteogenomics of Prostate Cancer MRI Visibility. Clinical Cancer Research 27, 3478–3490 (2021).

42. Vanaja DK, et al. Hypermethylation of genes for diagnosis and risk stratification of prostate cancer. Cancer investigation 27, 549–560 (2009).

43. Sivoňová MK, et al. Differential profiling of prostate tumors versus benign prostatic tissues by using a 2DE-MALDI-TOF-based proteomic approach. Neoplasma 68, 154–164 (2021).

44. Chen C, et al. Bioinformatics analysis of differentially expressed proteins in prostate cancer based on proteomics data. Onco Targets Ther 9, 1545–1557 (2016).

45. Wu J-P, et al. Intensity of stromal changes is associated with tumor relapse in clinically advanced prostate cancer after castration therapy. Asian Journal of Andrology 16, 710 (2014).

46. Zhao HB, et al. [Bioinformatics-based identification of the key genes associated with prostate cancer]. Zhonghua Nan Ke Xue 27, 489–498 (2021).

47. Cox DRJJotRSSSB. Regression models and life-tables. 34, 187–202 (1972).

48. Harrell Jr FE, Lee KL, Mark DBJSim. Multivariable prognostic models: issues in developing models, evaluating assumptions and adequacy, and measuring and reducing errors. 15, 361–387 (1996).

49. Ishwaran H, Kogalur UB, Blackstone EH, Lauer MSJTaoas. Random survival forests. 2, 841–860 (2008).

50. Jin HJ, Kim J, Yu J. Androgen receptor genomic regulation. Transl Androl Urol 2, 157–177 (2013).

51. Butler LM, et al. Lipids and cancer: Emerging roles in pathogenesis, diagnosis and therapeutic intervention. Advanced drug delivery reviews 159, 245–293 (2020).

52. Zadra G, et al. Inhibition of de novo lipogenesis targets androgen receptor signaling in castrationresistant prostate cancer. Proceedings of the National Academy of Sciences 116, 631–640 (2019).

53. Zhang A, Zhang J, Plymate S, Mostaghel EA. Classical and non-classical roles for pre-receptor control of DHT metabolism in prostate cancer progression. Hormones and Cancer 7, 104–113 (2016).

54. Sinha A, et al. The proteogenomic landscape of curable prostate cancer. Cancer Cell 35, 414–427. e416 (2019).

55. Chandrashekar DS, et al. UALCAN: a portal for facilitating tumor subgroup gene expression and survival analyses. Neoplasia 19, 649–658 (2017).

56. Intasqui P, Bertolla RP, Sadi MV. Prostate cancer proteomics: clinically useful protein biomarkers and future perspectives. Expert review of proteomics 15, 65–79 (2018).

57. Jin X, et al. Urine Exosomal AMACR Is a Novel Biomarker for Prostate Cancer Detection at Initial Biopsy. Frontiers in oncology 12, 904315 (2022).

58. Biswas S, Talukdar M. DIAGNOSTIC UTILITY OF AMACR EXPRESSION TO DIFFERENTIATE PROSTATE CARCINOMA FROM BENIGN HYPERPLASIA OF PROSTATE--A HOSPITAL BASED CROSS-SECTIONAL STUDY. Journal of Evolution of Medical and Dental Sciences 8, NANA (2019).

59. Alinezhad S, et al. Global expression of AMACR transcripts predicts risk for prostate cancer–a systematic comparison of AMACR protein and mRNA expression in cancerous and noncancerous prostate. BMC urology 16, 1–10 (2016).

60. Liu Q, et al. Malate dehydrogenase 2 confers docetaxel resistance via regulations of JNK signaling and oxidative metabolism. The Prostate 73, 1028–1037 (2013).

61. Eyvazi S, Farajnia S, Dastmalchi S, Kanipour F, Zarredar H, Bandehpour M. Antibody based EpCAM targeted therapy of cancer, review and update. Current Cancer Drug Targets 18, 857–868 (2018).

62. Hu Y, Wu Q, Gao J, Zhang Y, Wang Y. A meta-analysis and The Cancer Genome Atlas data of prostate cancer risk and prognosis using epithelial cell adhesion molecule (EpCAM) expression. BMC urology 19, 1–8 (2019).

63. Uhlen M, et al. Towards a knowledge-based Human Protein Atlas. Nat Biotechnol 28, 1248–1250 (2010).

64. Uhlen M, et al. A pathology atlas of the human cancer transcriptome. Science 357, (2017).

65. Liu H, Li L, Fan Y, Lu Y, Zhu C, Xia W. Construction of Potential Gene Expression and Regulation Networks in Prostate Cancer Using Bioinformatics Tools. Oxidative medicine and cellular longevity 2021, (2021).

66. Tinholt M, et al. Coagulation factor V is a marker of tumor-infiltrating immune cells in breast cancer. Oncoimmunology 9, 1824644 (2020).

67. Fuhrmann DC, Wittig I, Brüne B. TMEM126B deficiency reduces mitochondrial SDH oxidation by LPS, attenuating HIF-1α stabilization and IL-1β expression. Redox biology 20, 204–216 (2019).

68. Fuhrmann DC, Wittig I, Dröse S, Schmid T, Dehne N, Brüne B. Degradation of the mitochondrial complex I assembly factor TMEM126B under chronic hypoxia. Cellular and Molecular Life Sciences 75, 3051–3067 (2018).

69. Pezzuto A, Carico E. Role of HIF-1 in cancer progression: novel insights. A review. Current molecular medicine 18, 343–351 (2018).

70. Li Y, et al. Hypoxia-inducible miR-182 enhances HIF1α signaling via targeting PHD2 and FIH1 in prostate cancer. Scientific reports 5, 1–13 (2015).

71. Cui Y, Han B, Zhang H, Liu H, Zhang F, Niu R. Identification of Metabolic-Associated Genes for the Prediction of Colon and Rectal Adenocarcinoma. OncoTargets and therapy 14, 2259 (2021).

72. Lehrer S, Rheinstein PH. EARS2 significantly coexpresses with PALB2 in breast and pancreatic cancer. Cancer Treatment and Research Communications 32, 100595 (2022).

73. Wokołorczyk D, et al. PALB2 mutations and prostate cancer risk and survival. British Journal of Cancer 125, 569–575 (2021).

74. Carreira S, et al. Biomarkers Associating with PARP Inhibitor Benefit in Prostate Cancer in the TOPARP-B TrialPredictive Biomarkers for PARP Inhibition in Prostate Cancer. Cancer discovery 11, 2812–2827 (2021).

75. Kimura H, et al. Prognostic significance of pathogenic variants in BRCA1, BRCA2, ATM and PALB2 genes in men undergoing hormonal therapy for advanced prostate cancer. British Journal of Cancer, 1–11 (2022).

76. Zhang L, et al. Five-gene signature associating with Gleason score serve as novel biomarkers for identifying early recurring events and contributing to early diagnosis for Prostate Adenocarcinoma. Journal of Cancer 12, 3626 (2021).

77. Van den Broeck T, et al. Biochemical recurrence in prostate cancer: the European Association of Urology prostate cancer guidelines panel recommendations. European urology focus 6, 231–234 (2020).

78. Gharzai LA, et al. Intermediate clinical endpoints for surrogacy in localised prostate cancer: an aggregate meta-analysis. The Lancet Oncology 22, 402–410 (2021).

79. Van den Broeck T, et al. Prognostic value of biochemical recurrence following treatment with curative intent for prostate cancer: a systematic review. European urology 75, 967–987 (2019).

80. Guo T, et al. Rapid mass spectrometric conversion of tissue biopsy samples into permanent quantitative digital proteome maps. Nat Med 21, 407–413 (2015).

81. Demichev V, Messner CB, Vernardis SI, Lilley KS, Ralser M. DIA-NN: neural networks and interference correction enable deep proteome coverage in high throughput. Nat Methods 17, 41–44 (2020).

82. Cox J, Hein MY, Luber CA, Paron I, Nagaraj N, Mann M. Accurate proteome-wide label-free quantification by delayed normalization and maximal peptide ratio extraction, termed MaxLFQ. Mol Cell Proteomics 13, 2513–2526 (2014).

83. Shannon P, et al. Cytoscape: a software environment for integrated models of biomolecular interaction networks. 13, 2498–2504 (2003).

