## SupplementaryInformation for "Proteomic-based stratification of intermediate-risk prostate cancer patients"

**Supplementary Table 1. Clinicopathological features of patients with prostate cancer (PCa).**

| | | |
|---|---|---|
| No. of patients | 278 (277 tumour and 278 benign) | |
| No. of patients with follow-up | 198 | |
| Median follow-up (months) | 59 | |
| Range (months) | 0-93 | |
| **Clinicopathological characteristics** | | |
| **Variable** | **n** | **%** |
| Age at diagnosis (median = 64 years, range = 41-83) | | |
| <64 | 132 | 47.6 |
| ≥ 64 | 145 | 52.3 |
| Gleason grade | | |
| GG1 | 15 | 5.4 |
| GG2 | 134 | 48.3 |
| GG3 | 70 | 25.2 |
| GG4 | 29 | 10.4 |
| GG5 | 29 | 10.4 |
| Tumour stage (pT) | | |
| pT1 | 159 | 57.4 |
| pT2 | 79 | 28.5 |
| Unknown | 39 | 14 |
| Surgical margin | | |
| Negative | 145 | 52.3 |
| Positive | 93 | 33.5 |
| Unknown | 39 | 14 |
| Tumour percentage | | |
| 40% | 1 | 0.36 |
| 50% | 13 | 4.7 |
| 60% | 28 | 21.6 |
| 70% | 78 | 25.2 |
| 80% | 136 | 28.8 |
| 90% | 21 | 32.4 |
| Biochemical recurrence events | | |
| Missing | 78 | 28.1 |
| No | 168 | 60.6 |
| Yes | 31 | 11.1 |

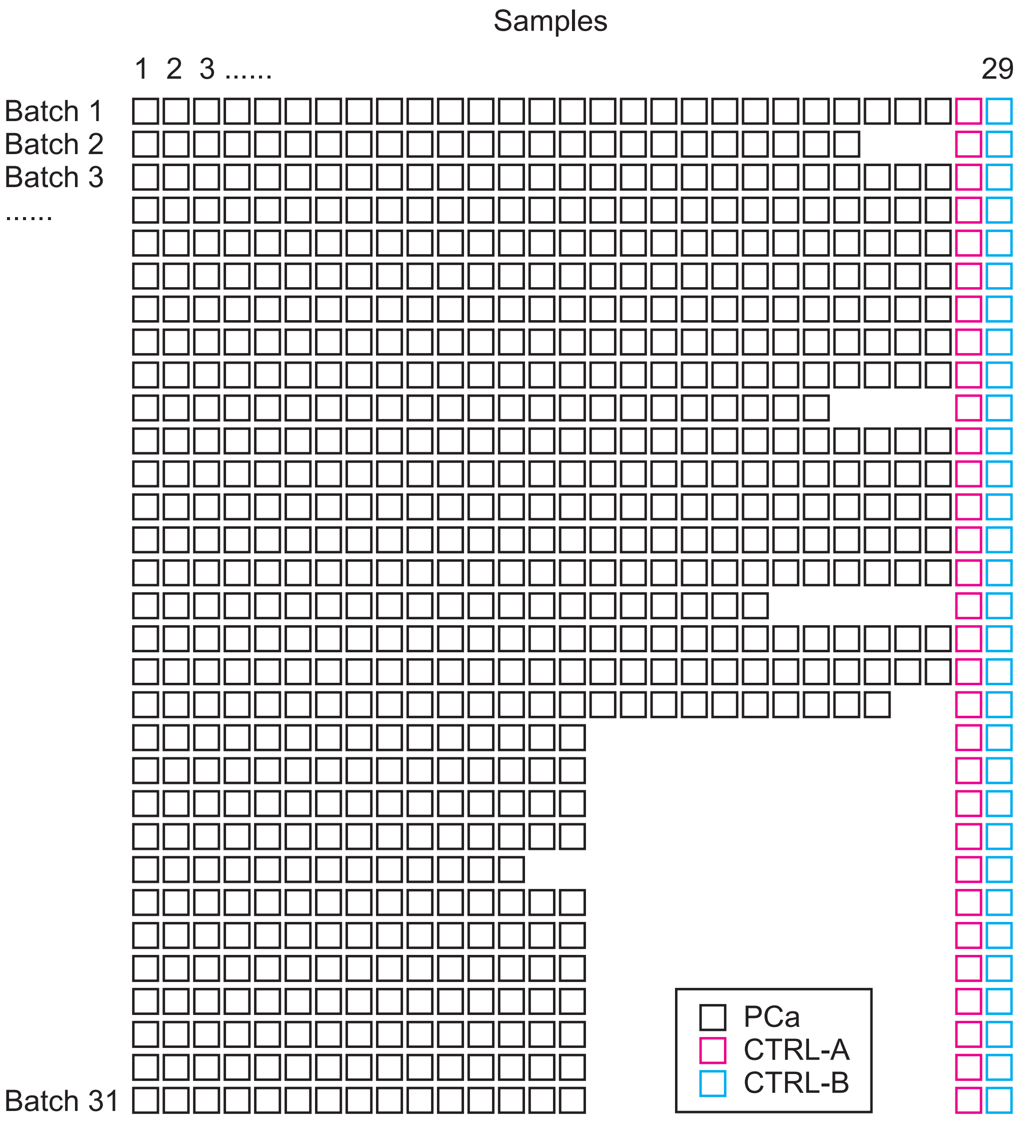


**Supplementary Figure 1. PPP1 study design.** Each row indicates a batch and each column indicates PCa tissue samples. Each batch contained between 15 and 29 PCa samples, one CTRL-A, and one CTRL-B.

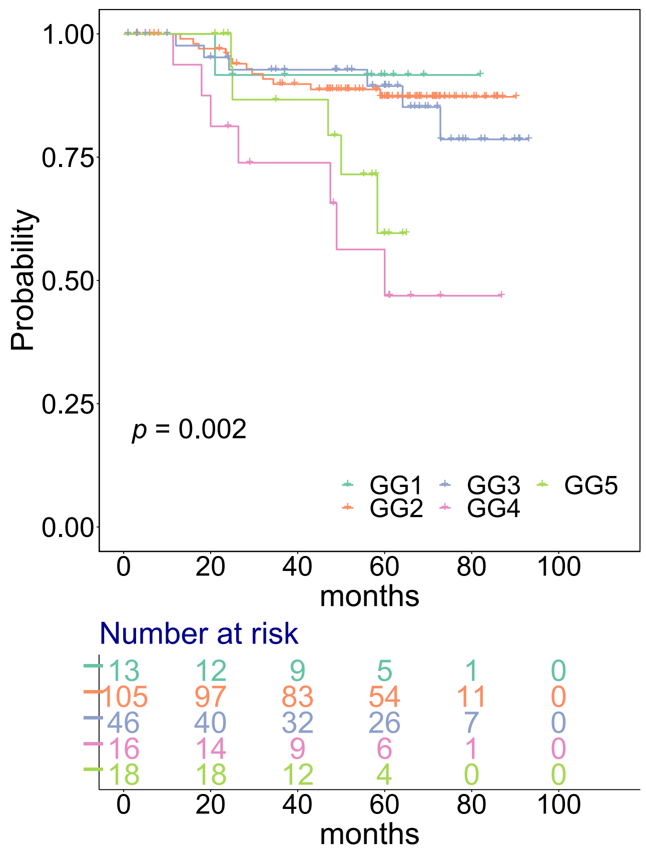


**Supplementary Figure 2. Kaplan-Meier (KM) curves for biochemical recurrence free survival (BCRFS) for different GGs.** Vertical lines illustrate patients who were censored at the time of their last clinical follow-up visit. The *p*-value shows significance of the difference between survival estimates evaluated by the log-rank test. Coloured values represent the number of patients in each group under risk.

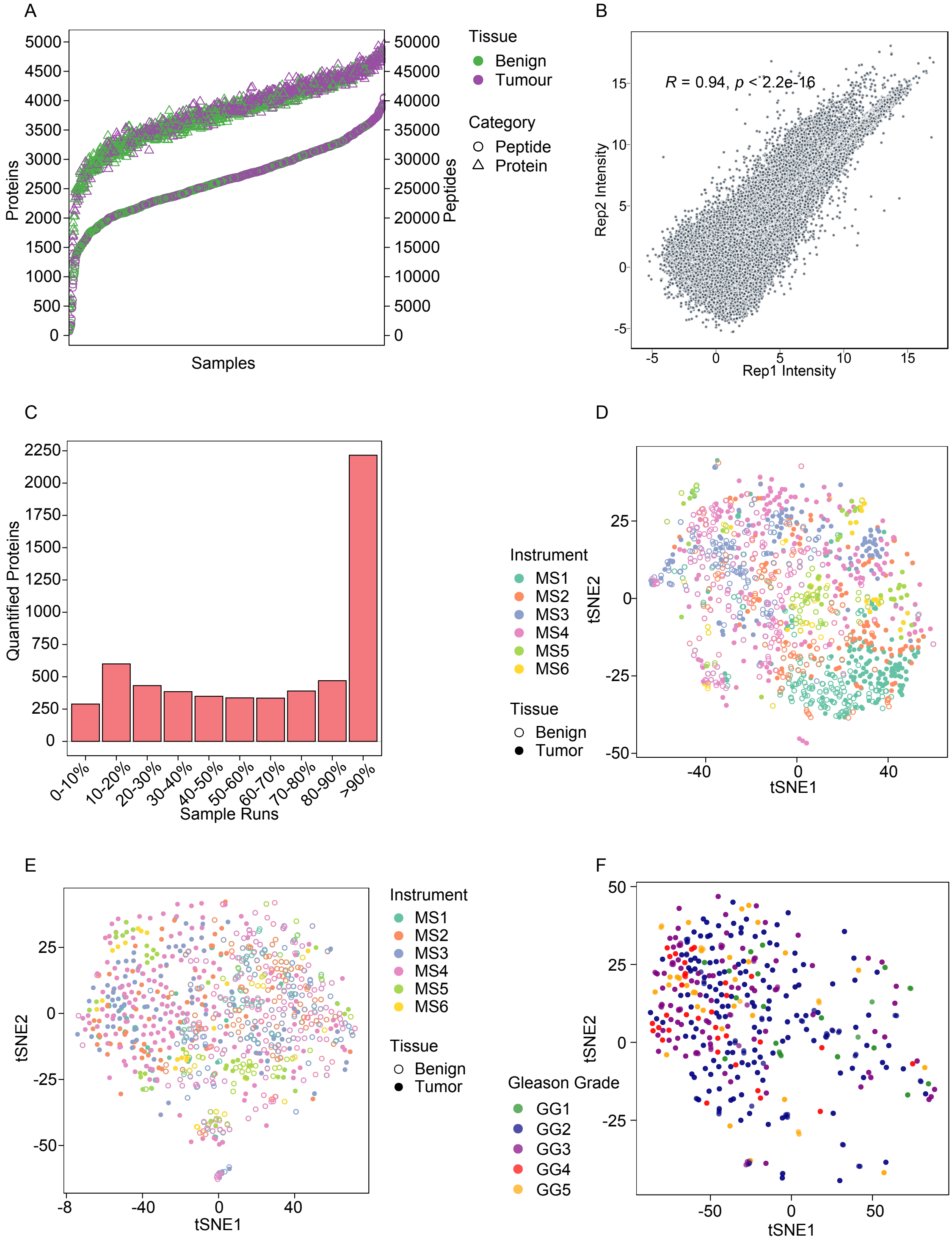


**Supplementary Figure 3. Overview of proteomics data. A.** Proteins and peptides quantified in tumour and benign samples (n = 1,348). A total of 53,713 peptides and 5,803 proteins are quantified. Purple colour shows the number of proteins and peptides quantified in tumour samples and green colour shows in benign samples. Compared to tumour samples, a smaller number of proteins are quantified in benign samples. **B.** The technical reproducibility of dataset shown by Pearson's r among replicates. **C.** Proteins quantified in different fractions of samples. Approximately 3,500 proteins are quantified in 50% or more samples. **D.** tSNE projection of proteomic data before batch correction. Samples analyzed using different mass spectrometers (MS1-6) are shown with different colors. Clusters of tumour and benign samples are shown in different shapes. **E.** tSNE projection of proteomic data after batch correction showing no batch effects. **F.** tSNE projection of tumour samples coloured by GG. No grouping can be observed.

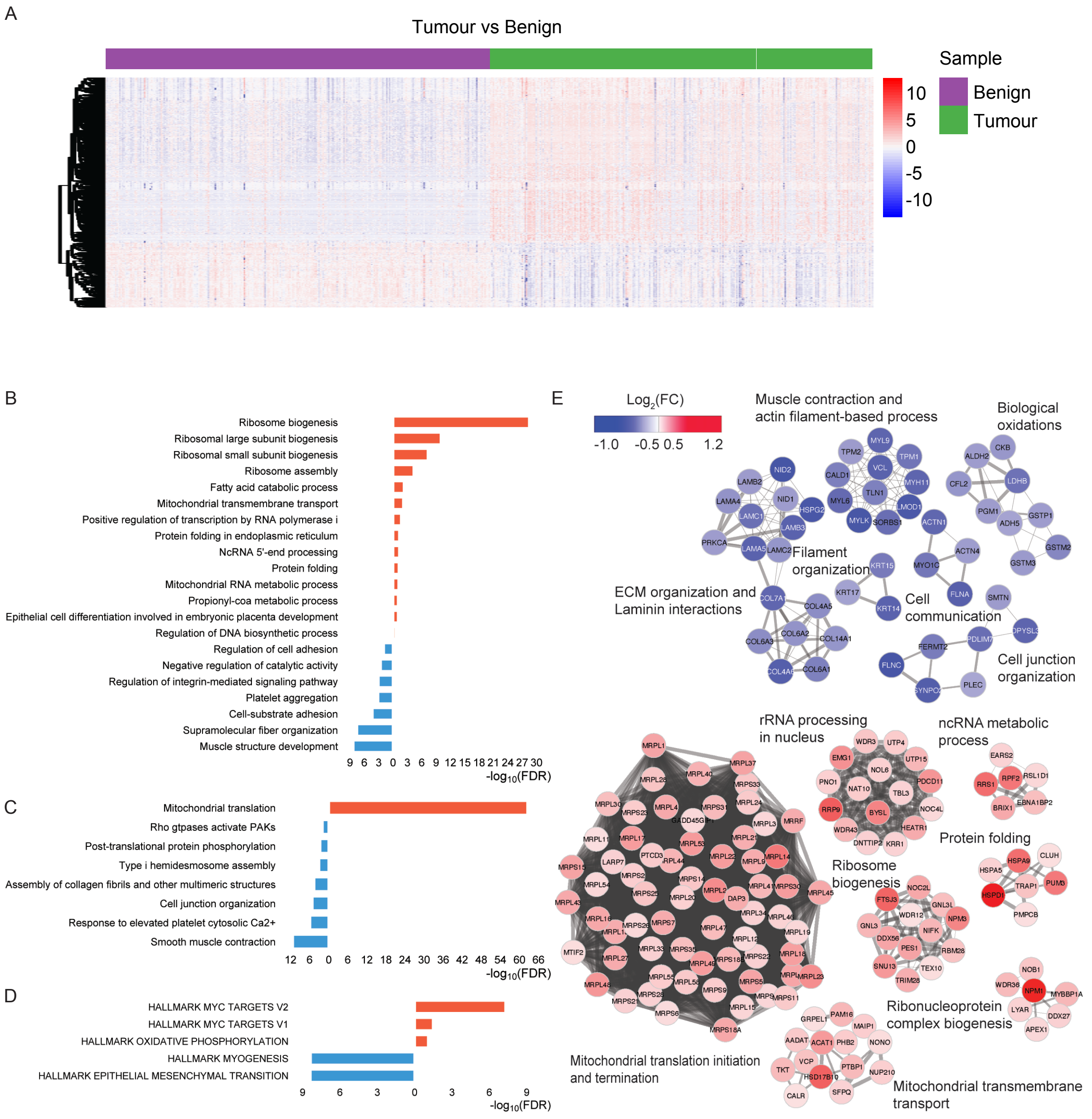


**Supplementary Figure 4. Dysregulated pathways in tumour samples. A.** Heatmap representation of the expression levels of differentially expressed proteins between tumour and benign samples shown in **Figure 1D**. Expression values are converted to z-scores. Samples are sorted according to tissue types (tumour vs. benign) on the x-axis whereas proteins are clustered on the y-axis. Biological pathways including GO biological processes (**B**), Reactome pathways (**C**) and hallmark gene sets (**D**) enriched for the tumor vs benign differentially expressed proteins**.** Red bars indicate pathways enriched in upregulated proteins and blue bars indicate pathways enriched in downregulated proteins. **E.** PPI network components were obtained using MCODE algorithm, showing the enriched biological processes and proteins. Upregulated and downregulated networks and associated proteins are coloured by FC. Upregulated proteins are coloured in red and downregulated in blue. The width of the edge (between nodes) indicates the strength of the connection. A functional description is provided beside each component.

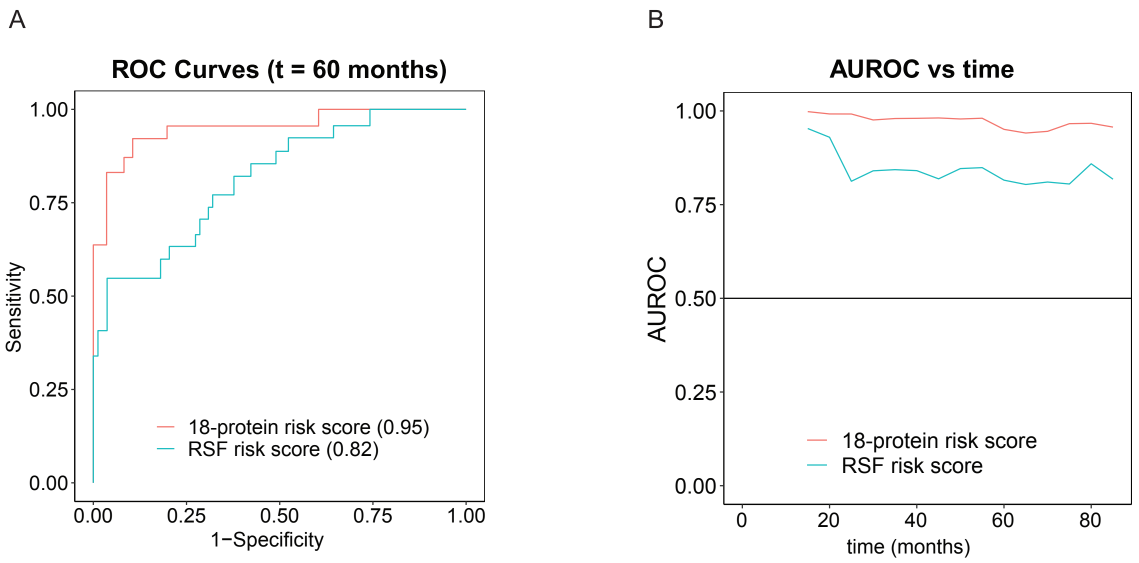


**Supplementary Figure 5. Survival Analysis of BCRFS. A.** ROC curves with respective AUROCs at five-year (60 months) follow up for the 18-protein risk score and RSF-based risk score **B.** Time-dependent AUROCs of the 18-protein risk score and RSF-based risk score.

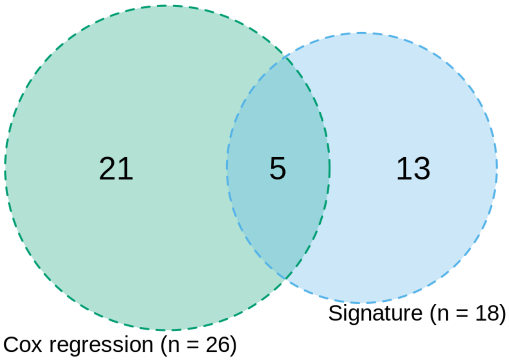


**Supplementary Figure 6. A set of unique proteins.** A set of 39 unique proteins was extracted by taking the union of 18 signature proteins and 26 proteins from a univariate Cox regression. Five proteins were found overlapping between the two sets.
